# Edible formulations of chicken egg derived IgY antibodies neutralize SarsCoV2 Omicron RBD binding to human ACE2

**DOI:** 10.1101/2022.02.21.481267

**Authors:** Kranti Meher, S. Sivakumar, Gopi Kadiyala, Subramanian Iyer, Subhramanyam Vangala, Satish Chandran, Uday Saxena

## Abstract

SarsCoV2 virus driven pandemic continues to surge propelled by new mutations such as seen in Omicron strain. Omicron is now rapidly becoming the dominant strain globally with more than 30 mutations in the spike protein. The mutations have resulted in Omicron strain escaping most of the neutralizing antibodies generated by the current set of approved vaccines and diluting the protection offered by the vaccines and therapeutic monoclonal antibodies.

This has necessitated the need for newer strategies to prevent this strain from spreading. Towards this unmet need we have developed chicken egg derived anti-RBD IgY antibodies that neutralize the binding of Omicron RBD to human ACE2. Furthermore, we have formulated the edible IgY as flavored beverages to allow for use as oral rinse and prevent the entry of Omicron in the oropharyngeal passage, a major access and accumulation point for this strain in humans.

## Introduction

The new SarsCoV2 virus Omicron strain has triggered another wave of COVID 19 pandemic globally. Omicron has more than 30 mutations in its spike protein that are believed to drive its higher transmission (1). In addition, this strain unlike the previous strains such as delta appears to reside mainly in the upper respiratory airways/oral cavity and not the lung (2). Therefore, strategies aimed at curbing this strain have to focus on this disparity between the various strains into account.

Another perhaps more worrisome aspect of Omicron strain is it escapes most of the neutralizing antibodies raised in response to currently used vaccines (3). Studies have shown that unlike Delta strain, the Omicron strain receptor binding domain is not neutralized in its binding to human ACE2 receptor by bulk of the antibodies produced in humans as well as some of the monoclonal antibody therapeutics (4). How then can we combat Omicron spread? A booster dose of the currently used vaccines has been approved by many countries in the hope that it may lead to higher titer of neutralizing antibodies but it remains to be seen if can help curb Omicron infections.

We have developed anti-RBD IgY antibodies from chicken eggs by immunizing the birds with RBD domain of SarsCov2 spike protein as a strategy to block the binding of virus RBD to human ACE2 receptors. Our intention is to use these IgY as a prophylactic against COVID 19. Chicken egg derived IgY have several advantages including being edible since they are considered as GRAS (generally regarded as safe). Importantly IgY are polyclonal in nature, which means to a given antigen, unlike monoclonal antibodies which target only a specific antigenic epitope, IgY have the ability to potential to hit multiple epitopes and greatly enhance the possibility of neutralizing activity. Such an approach is especially relevant in situations such as Omicron infections where the strain seems to “escape” the naturally produced antibodies.

We demonstrate here that our IgY raised against native RBD:

1. Completely neutralize the binding of Native, Delta and Omicron strain RBD to human ACE2
2. Formulations of IgY as flavored beverages retain the neutralizing activities
3. Such formulations are useful as prophylactic oral rinse against potential exposure especially to Omicron the dominant strain worldwide currently

## Methods

1. Preparation of anti-RBD IgY Anti-RBD IgY used here were prepared exactly as described before (5) and used at 5 mg/ml or otherwise indicated.
2. Dot blot protocol For dot blot studies shown here we used nitrocellulose membrane from 3M. RBD and biotinylated ACE2 were procured and used as described before. Briefly RBD was spotted at desired quantity in 10 ul of buffer onto the membrane, followed by washing with PBS wash buffer and PBS blocking buffer containing 3% FBS. After incubation of biotinylated human ACE2 and incubation, color generating substrate was added and absorbance read at OD540. Image J software analysis was used to quantitate the dot blot images. Typically, 0.25 ug/ml of RBDs and 0.25 or 0.5 ug/ml of biotinylated human ACE2 were used.
3. ELISA protocol The ELISA protocol used here was essentially the same as described before (5).
4. The whole egg powder from eggs of chickens of RBD immunized birds were used for the flavored beverage formulations. Typically, the IgY content in the formulations was 5 ug/ml unless otherwise indicated.

## Results

### 1. Neutralization of Omicron RBD binding to human ACE2 by IgY

We first tested if purified anti-RBD IgY raised against native RBD is able to neutralize the binding of biotinylated human ACE2 to Native, Delta and Omicron RBD coated on to nitrocellulose membrane in a dot blot format. Nitrocellulose strips were coated with respective RBD’s at 0.25 ug and binding of biotinylated human ACE-2 (0.25ug) in presence or absence of excess anti-RBD IgY (50 ug) was examined. As seen in figure 1, the binding to human ACE2 to all three RBD’s was blocked almost completely by anti-RBD IgY (figure 2 shows quantitation by image J software analysis) suggesting that strong neutralising activity. More significantly the neutralising activity of the IgY against Omicron RBD opens up possibility of using the IgY as a defence against this strain.

**Figure 1.**
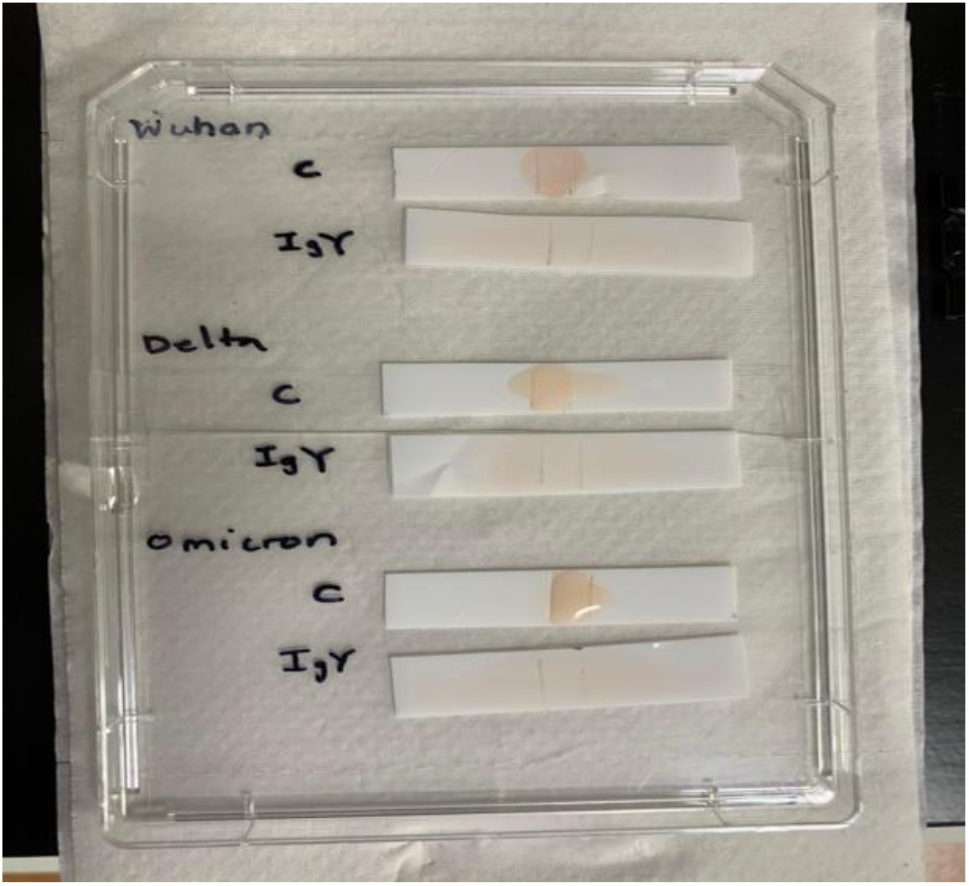
Anti-RBD IgY completely block ACE-2 interaction with Native, Delta and Omicron RBD

**Figure 2:**
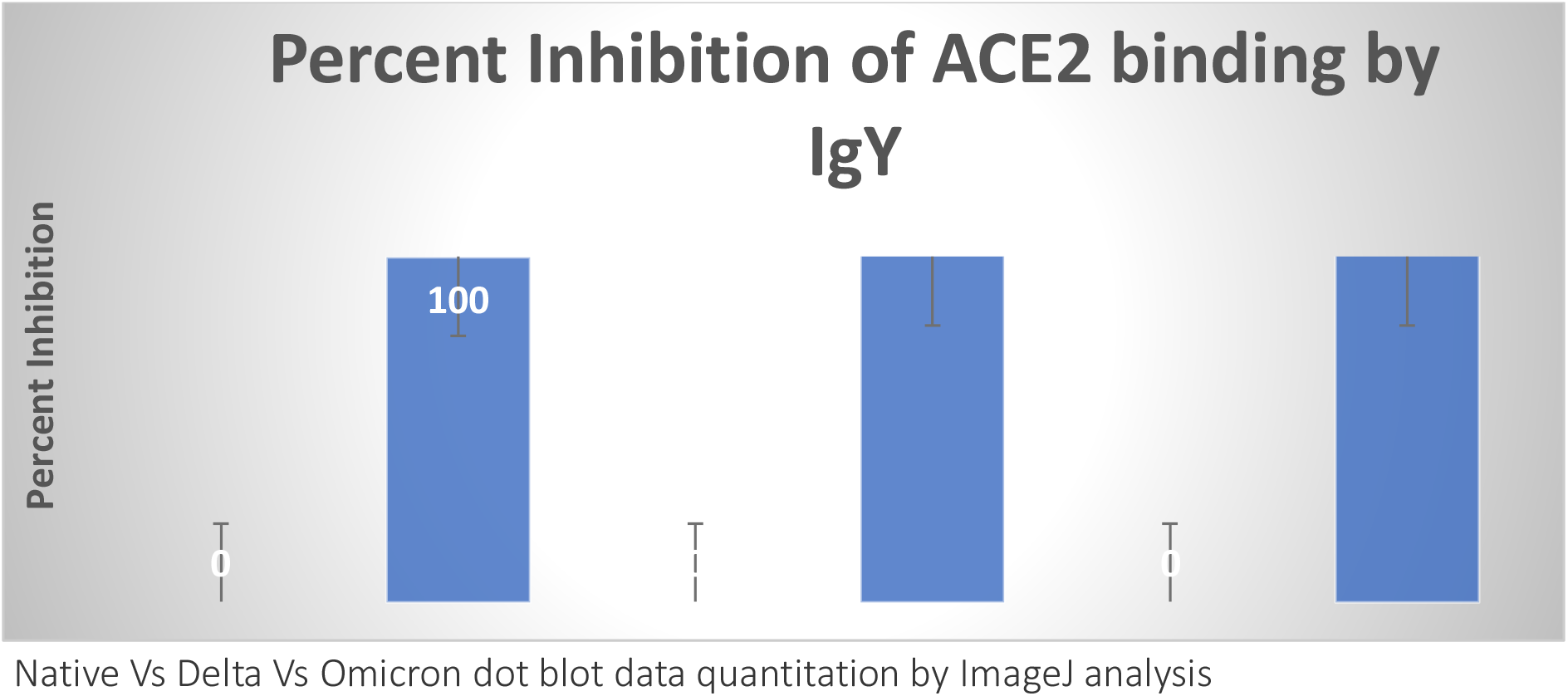
*P<0.05 for all data points Student’s t-tests were performed to compare between the control (no IgY added) versus presence of IgY (50 ug). RBDs were used at 0.25ug/ml.

### 2. Flavored beverages containing anti-RBD IgY retain neutralizing activity

To further test the feasibility of using the anti-RBD IgY as edible defence against binding of viral RBD to ACE2 receptors, we formulated the IgY in flavoured beverages and first tested against Native RBD. If successful in neutralizing, the beverages could be used as oral rinse prophylactic against virus entry through the oropharyngeal cavity.

As shown in figure 3, peppermint, grape and lemon flavoured beverages containing retained their neutralising activity (> 85% compared to unformulated IgY) suggesting that the IgY could be formulated as palatable beverage flavours and still retain neutralising activity.

**Figure 3:**
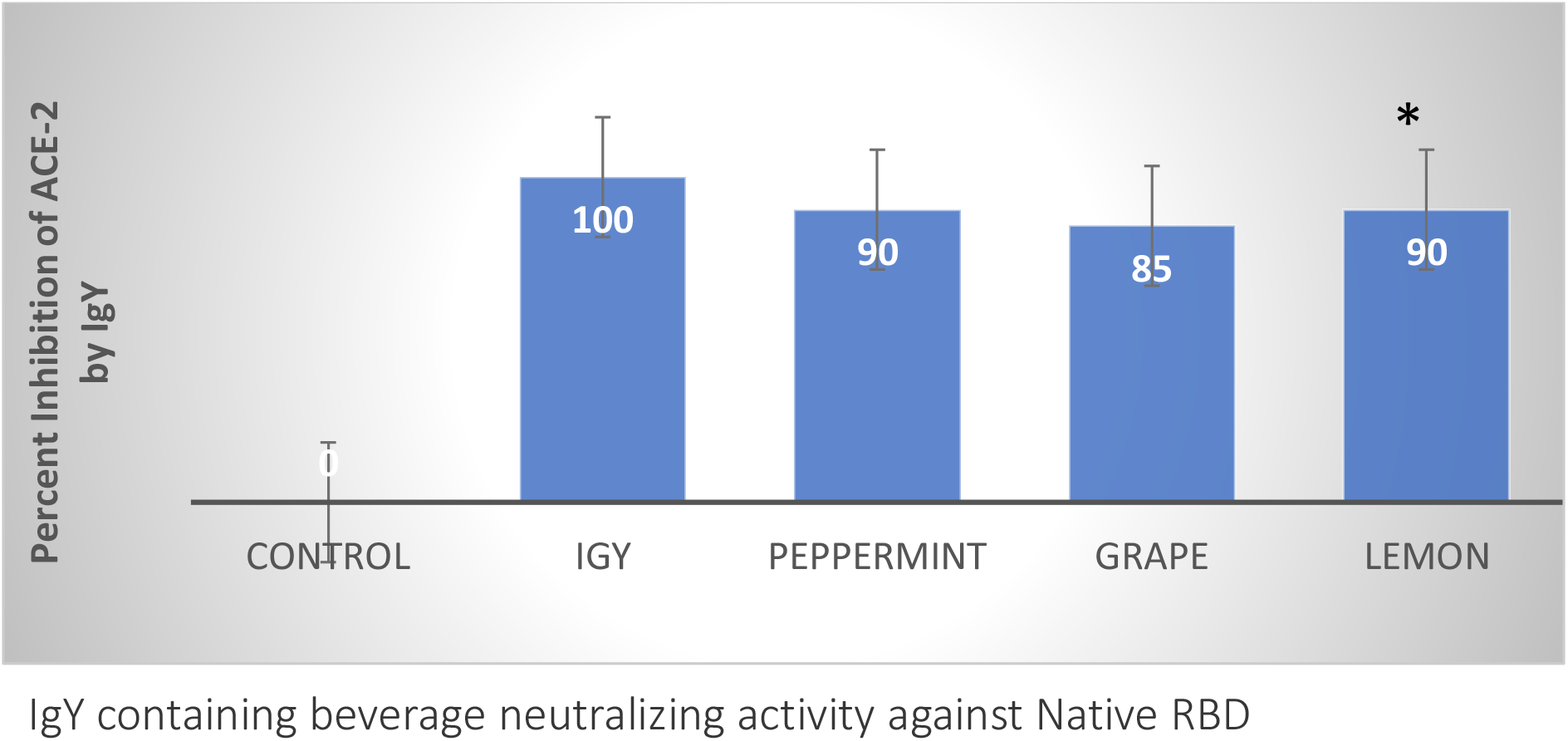
All the data points had *P<0.05 or greater. Student’s t-tests were performed to compare between the control (on IgY) and different flavor beverages or pure IgY.

Encouraged by these data we further tested additional flavoured beverages neutralising activities against Native and Delta strain RBD. Specifically we tested chocolate, strawberry and raspberry flavoured beverages containing anti-RBD IgY. Figure 4 shows that these flavours also show neutralising activity albeit moderately less than the previously tested flours. These data suggest 1) neutralising activity is retained in a variety of flours and 2) the beverages are active against Native and Delta strain RBD suggesting a broad range of neutralising activities.

**Figure 4:**
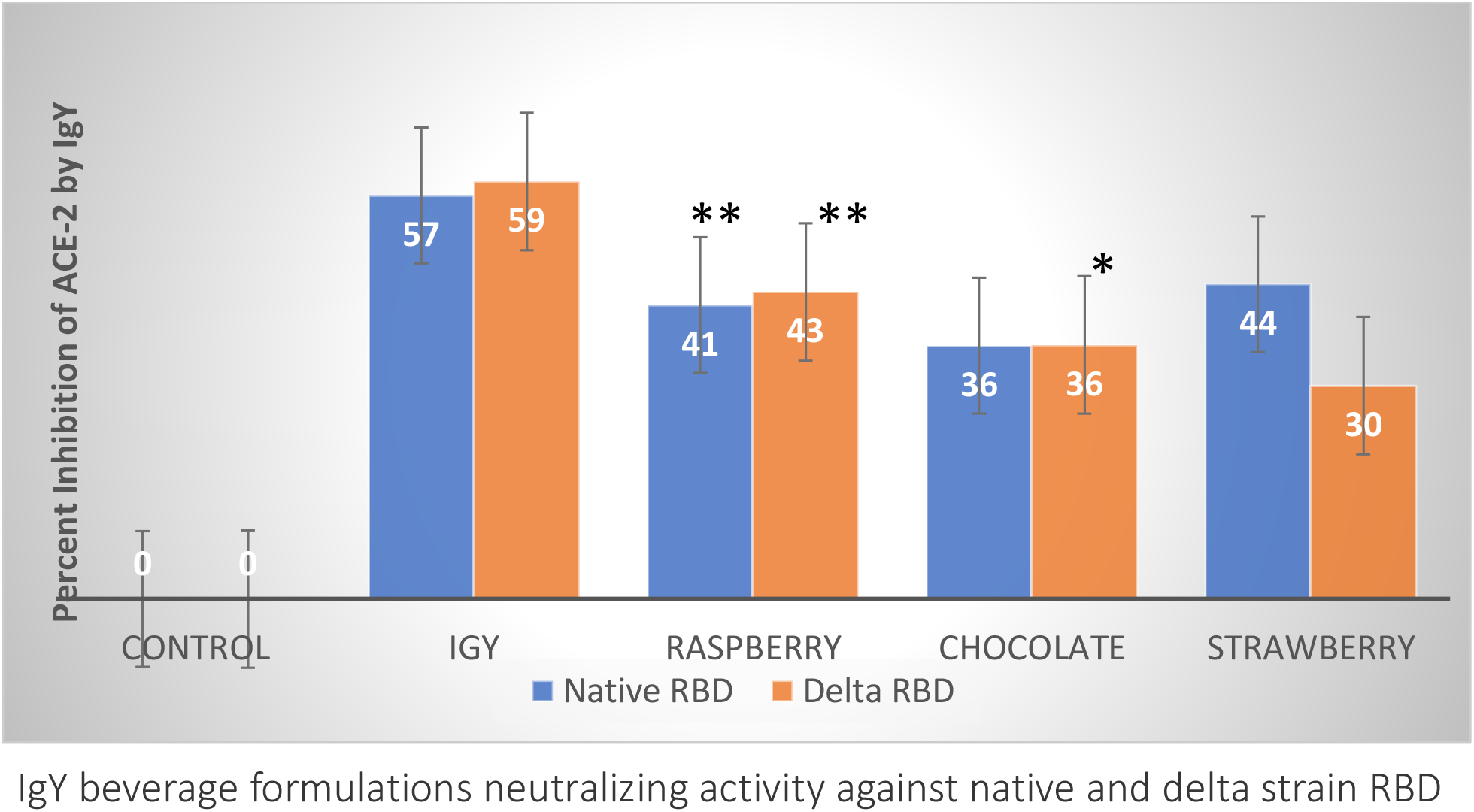
** P<0.01; *P<0.05. Student’s t-tests were performed to compare between the control and pure IgY (50ug) or beverages

We were most interested in testing anti-RBD containing flavoured beverages for neutralising activity against Omicron RBD. The impetus for this is that Omicron is currently the dominant strain globally, appears to reside more in oropharyngeal and upper respiratory airways and there is no known prophylactic anti-dote against it. Figure 5 shows that a variety of flavoured beverages are highly effective in neutralizing the binding of ACE2 to Omicron RBD with raspberry, peppermint and grape being most potent followed by chocolate strawberry and lemon flavours. These data strongly support the idea of us being able to use flavoured beverage formulated anti-RBD IgY mouth wash as an prophylactic protection against Omicron strain. Figure 6 presents a summary of neutralizing activity of all the flavours tested and shows that raspberry, peppermint and strawberry possess the best activities against Omicron RBD.

**Figure 5.**
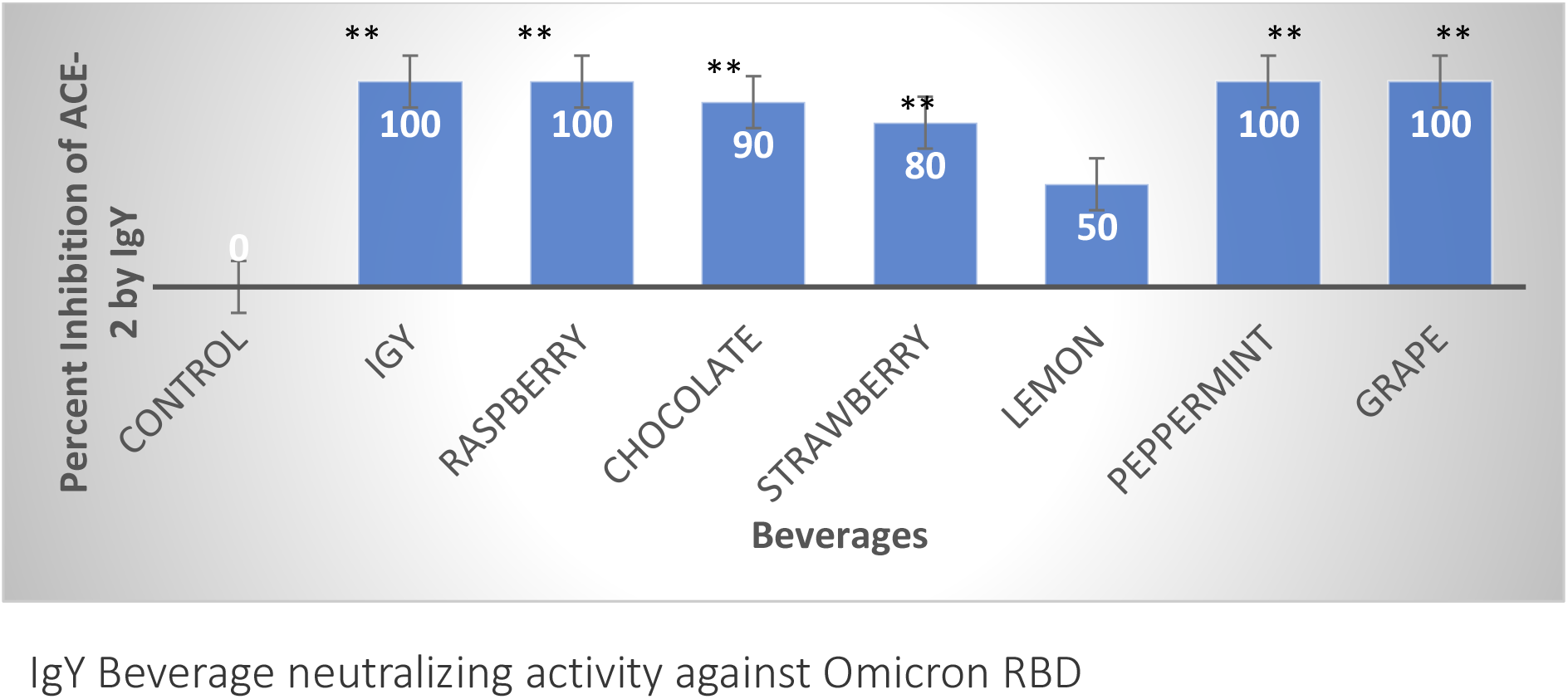
** P<0.01; *P<0.05. Student’s t-tests were performed to compare between the control and various flavors of beverages or pure IgY (50ug).

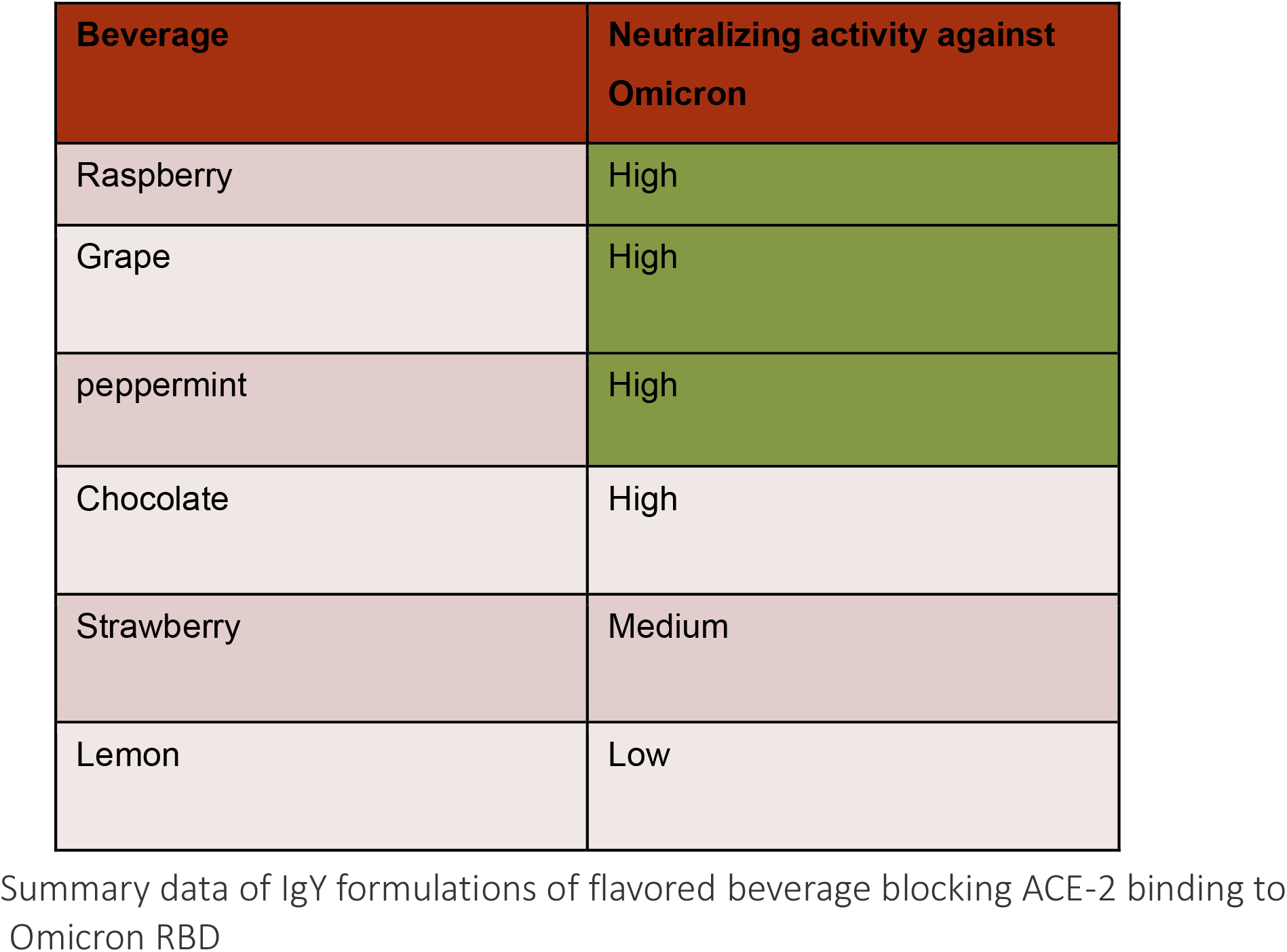

### 3. Binding of IgY to Omicron RBD

The neutralizing activity of our anti-RBD IgY could be either due to direct interaction between the anti-RBD IgY and the RBDs or due to other unknown mechanism of action. To test if there is binding of the IgY to RBDs we performed a direct ELISA assay where the RBD were coated onto the plate and the binding of our anti-RBD IGY assessed (using Absorbance Units, at OD 540) with HRP labelled anti-IgY for detection. A comparison of IgY binding to Native, Delta and Omicron RBDs was made.

Figure 7 shows the Absorbance Units profile of IgY binding to all three RBDs at a various amounts of RBD on the plate and fixed amount of IgY added. Firstly, it is obvious from the data that IgY binds to all three RBD’s although the degree of binding varies. Secondly the highlighted boxes in green show the AU_50_ (concentration of _IgY_ at which 50% percent of maximal Absorbance Units used as a measure of binding capability) shows that the AU_50_ of IgY binding to native RBD to be between 0.2-0.4 ug, for Omicron RBD it was 0.1 ug and for Delta was 0.1-0.2 ug suggesting they IgY binding to all three RBDs.

**Figure 7.**
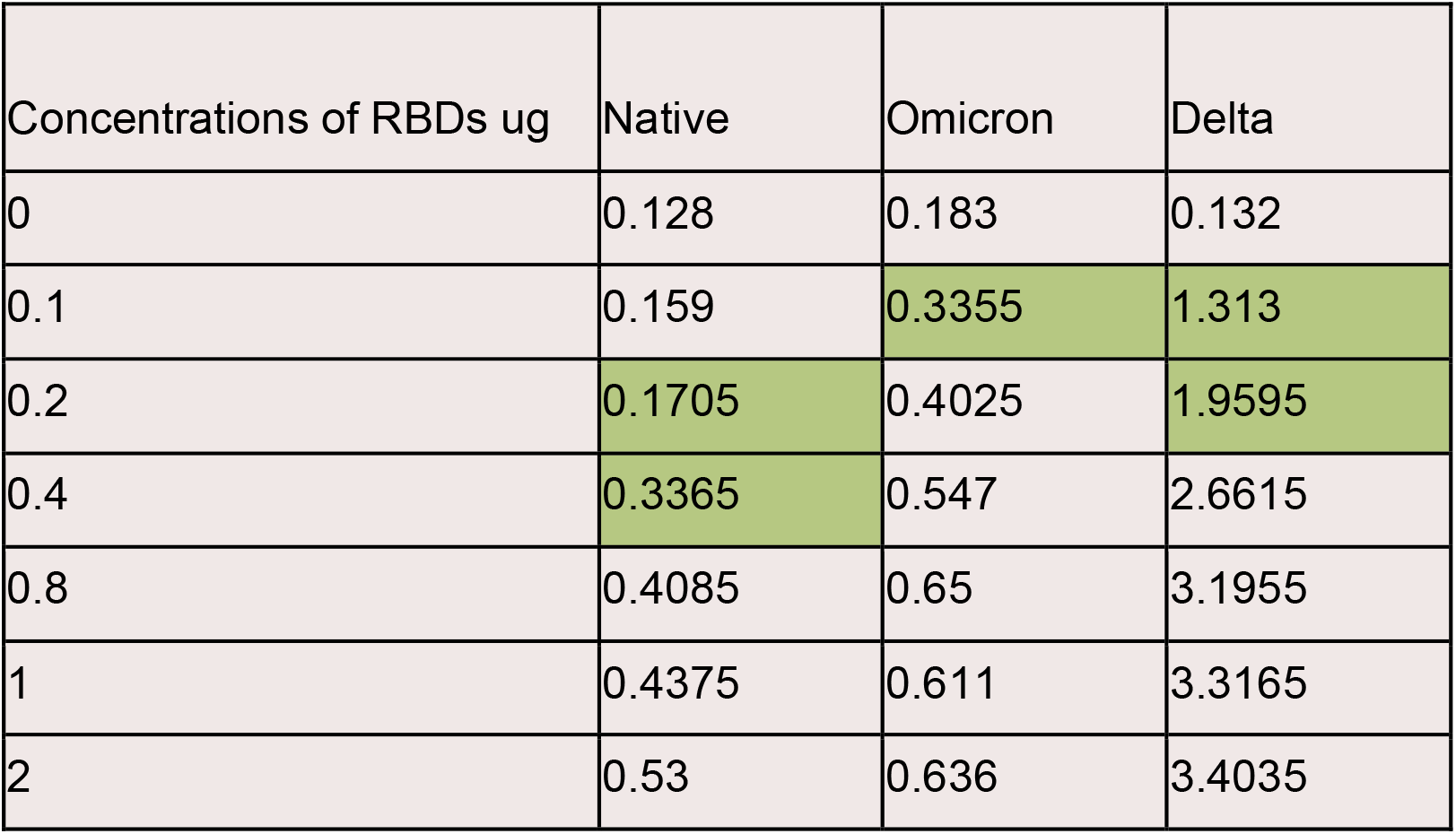
Binding of IgY to RBDs Absorbance Units OD 540

Despite similar AU_50_ for all three RBDs, in terms of total binding of IgY to Delta RBD was almost 5 times higher potentially suggesting higher binding capacity perhaps due to recognition of more epitopes of Delta. Figure 8 shows the same binding data in a graphical form.

**Figure 8:**
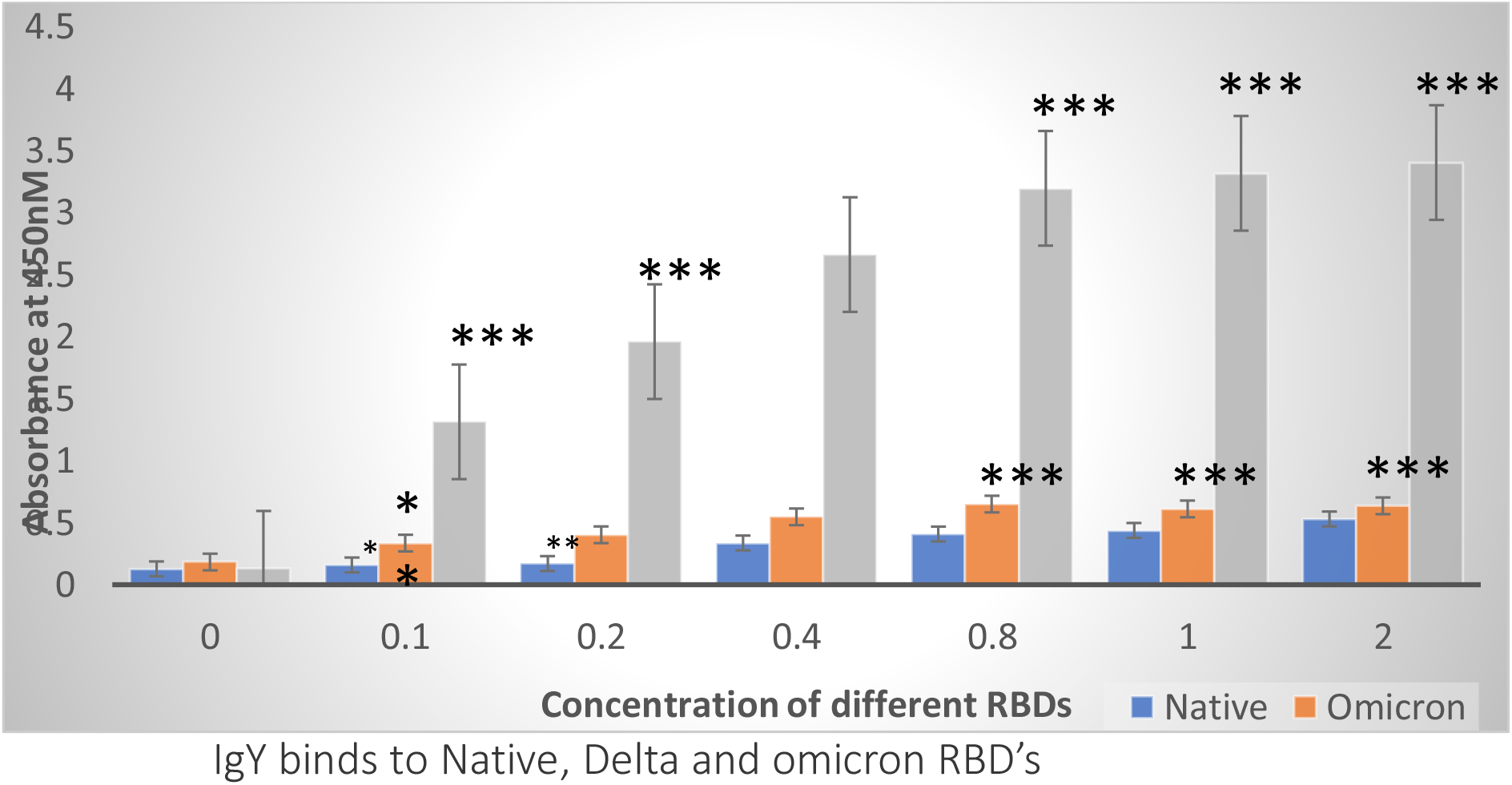
Dose response study. Different concentrations of RBDs against 2ug/mL of Anti-RBD IgY *** P<0.001; ** P<0.01; *P<0.05. Student’s t-tests were performed to compare between the control and different

To further explore the binding of IgY to the RBD’s we performed another ELISA wherein, the amount of RBD on the plate was fixed and interrogated with increasing amounts of anti-RBD IgY (0-8 ug). Figure 9 presents the data in terms of AU units and analysis of AU_50_ shows the value for Native RBD to be between 2-4 ug, for Omicron at 2 ug and for Delta to be between 0.4-0.8 ug. Figure 10 shows the same in a graphical format.

**Figure 9.**
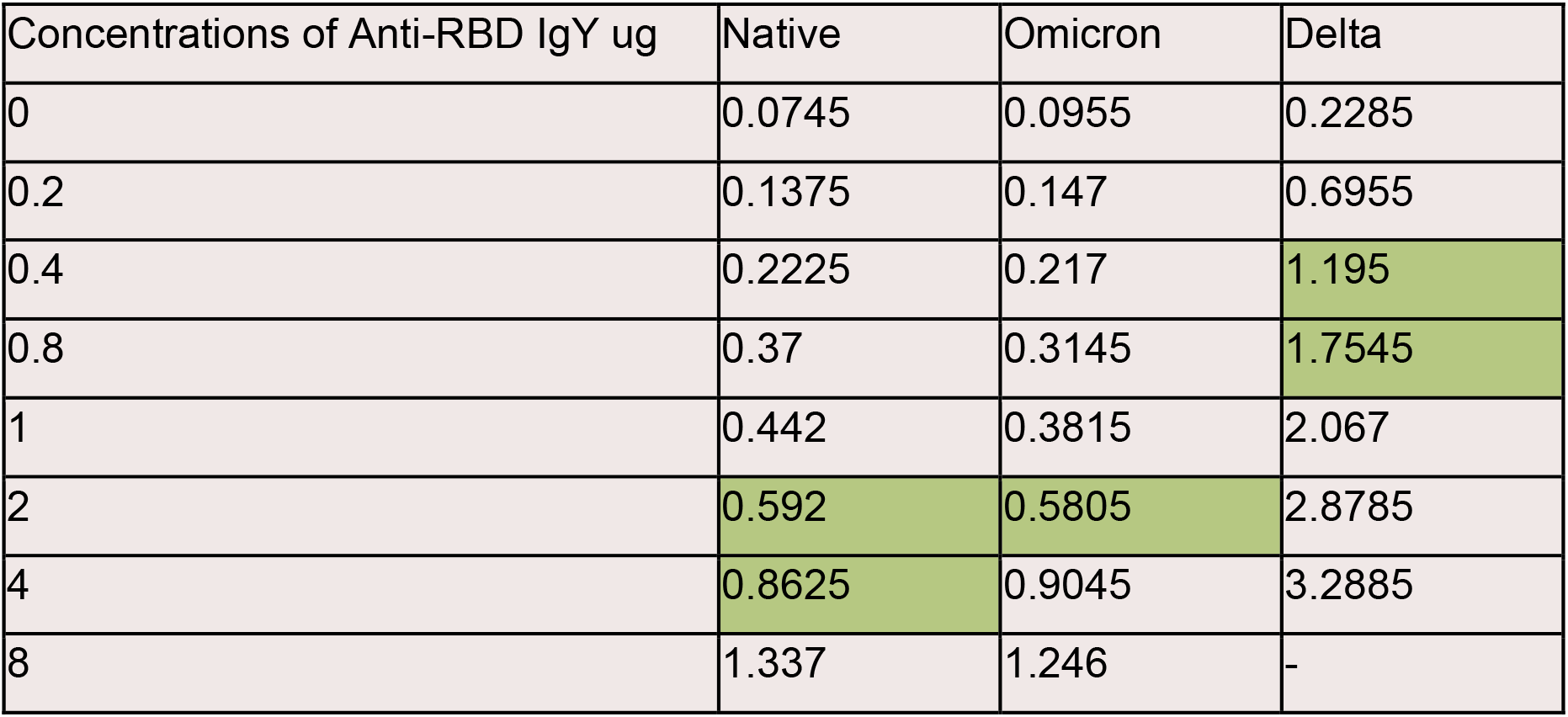
Binding of IgY to the three RBDs

**Figure 10:**
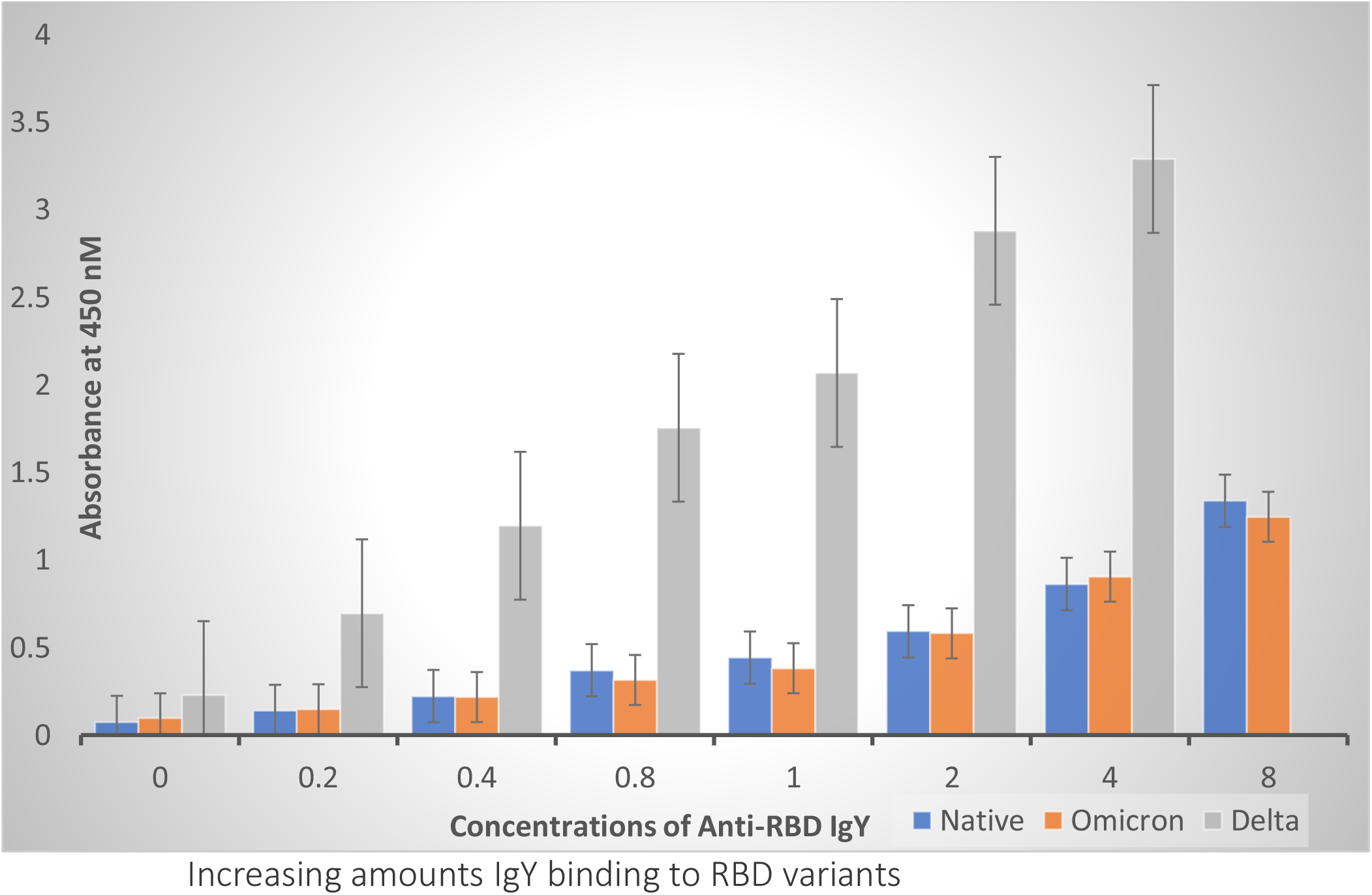
Different concentrations of Anti-RBD IgY binding to RBDs. All of the data points were at value of P<0.01 or better by Student’s t-tests compared between the control to binding various amounts of IgY

Collectively these ELISA data suggest that

1. Anti-RBD IgY binds to all three RBD’s
2. AU_50_ values in both experiments suggest the rank order to be Delta RBD> Omicron RBD > Native RBD
3. The binding of anti-RBD IgY to all three RBDs may be due to the polyclonal nature of IgY

## Discussion

The experiments presented here were designed to probe a) do anti-RBD IgY neutralize the binding of RBDs to human ACE2 especially Omicron? and b) Do the IgY bind to RBD?

Our data demonstrate complete neutralization of binding to all three RBDs to ACE2 by the IGY and also direct binding of IgY to RBDs as a potential mechanism by which this neutralization may occur.

We are most enthusiastic about the neutralization of Omicron strain RBD. The reasons for this are following

1. Currently there is no known remedy to counter the entry and binding of Omicron strain thru the oropharyngeal and upper respiratory airways, a preferred residence destination for this strain unlike Native and Delta strains which are able to effectively penetrate deep lung tissue
2. The ability to formulate the IgY in flavored beverages and retain neutralizing activity offers the possibility of using the beverages as an oral mouth wash or drink, made possible by the fact that egg IgY are considered safe for human consumption.

We envision a scenario where these beverages can be used as prophylactic in all comers, children, elderly, un-vaccinated and vaccinated. The reason for this proposed use is that recent data which shows that currently used vaccines do not illicit strong enough neutralizing response against Omicron, leaving even the vaccinated vulnerable. The formulations are currently in development for human use.

## Acknowledgements

We gratefully acknowledge the support in the experiments from Arpitha Reddy and Saranya K.

## References

1. The Lancet Comment| Volume 398, ISSUE 10317, P2126–2128, December 11, 2021, Omicron SARS-CoV-2 variant: a new chapter in the COVID-19 pandemic, SS Abdool Karim, Q Abdool Karim

2. Nature NEWS, 05 January 2022, Correction 06 January 2022. Omicron’s feeble attack on the lungs could make it less dangerous. Max Kozlov

3. BioRxiv preprint doi: https://doi.org/10.1101/2021.12.03.471045; Omicron-B.1.1.529 leads to widespread escape from neutralizing antibody responses, Wanwisa Dejnirattisai et al

4. Nature NEWS 21 December 2021. Omicron overpowers key COVID antibody treatments in early tests. Max Kozlov

5. BioRxiv preprint doi: https://doi.org/10.1101/2021.10.19.464951, Preparation of ingestible antibodies to neutralize the binding of SarsCoV2 RBD (receptor binding domain) to human ACE2 Receptor. Gopi Kadiyala et al

